# Ironing out the role of Nrf2 in cardiac iron metabolism during myocardial infarction

**DOI:** 10.1101/2024.09.25.615071

**Authors:** Deepthy Jayakumar, Kishore Kumar S. Narasimhan, Abinayaa Rajkumar, Gokul Prasanth Panchalingam, Navvi Chandrasekar, Varsha C. Ravikumar, Kalaiselvi Periandavan

**Affiliations:** Department of Medical Biochemistry, Dr. ALM Post Graduate Institute for Basic Medical Sciences, University of Madras, Chennai, 600113, Tamil Nadu, India; Department of Psychiatry and Behavioral Sciences, School of Medicine, Texas A&M University College Station, Texas, USA; Department of Pathology, Dr. ALM Post Graduate Institute for Basic Medical Sciences, University of Madras, Chennai, 600113, Tamil Nadu, India

**Keywords:** Ferroportin, Ferritinophagy, Hepcidin, Iron, Myocardial Infarction, Nrf2

## Abstract

**Background and Purpose:** Iron plays a crucial role in maintaining cardiac health. However, existing research has focused on understanding how cardiac cells regulates intercellular iron levels through their own cell-autonomous cardiac hepcidin/ferroportin axis. In Addition, several studies have explored the mechanisms linking cardiac dysfunction with iron imbalance. Recent insights also emphasize the importance of Nrf2, a key transcriptional regulator that not only counteracts iron-mediated oxidative stress, but also governs several genes involved in iron metabolism. Consequently, the Nrf2/hepcidin/ferroportin axis is emerging as a central hub connecting cardiac iron metabolism with redox alterations. However, the precise mechanisms linking these components remain elusive. This study aims to elucidate how disruptions in the Nrf2/hepcidin/ferroportin axis contribute to the altered iron metabolism in Myocardial infarction (MI).

**Experimental Approach:** MI was induced in adult Wistar rats by subcutaneous administration of isoproterenol (ISO; 85 mg/kg body weight) for two days. H9c2 cardiomyoblasts were differentiated into cardiomyocytes using all-trans-retinoic acid (ATRA, 2.5μM for 5-days) and subjected to hypoxic stress using CoCl_2_ (100μM). *In vitro* pharmacological suppression of Nrf2 was performed using brusatol (50nM).

**Key Results:** Morphological examination revealed maladaptive remodeling, and histopathological analysis demonstrated disoriented myofibrils with intense neutrophil infiltration and necrotic impressions in MI-affected animals. Furthermore, elevated levels of labile redox-active iron and inflammatory markers were observed in serum of ISO induced animals. qPCR & Western blot analysis indicated an increase in HIF-1α and hepcidin levels, and downregulation of FTH levels in MI-induced animals, with no significant changes observed in FPN-1. The transcriptional activity of Nrf2 is enhanced in the MI-heart. Moreover, increased levels of NCOA4, beclin-1, and LC3-II/LC3-I, along with decreased p62, suggest enhanced ferritinophagy in MI-induced hearts. Nrf2 was pharmacologically suppressed in differentiated H9c2 cardiomyocytes to explore its potential role in MI pathophysiology. Remarkably, this inhibition rescued CoCl_2_-induced hypoxic stress, as evidenced by the decreased ferritinophagy and apoptotic cell death.

**Conclusion and Implications:** Augmented Nrf2-transcriptional activity disrupts iron metabolism through the hepcidin/ferroportin axis, leading to iron sequestration and promoting ferritinophagy within cardiomyocytes, thereby exacerbating MI.

## 1. INTRODUCTION

The heart possesses a distinctive iron homeostasis profile owing to its substantial energy requirements, necessitating an increased demand for iron (Jayakumar et al., 2022). As a result, the regulation of oxygen levels and iron metabolism intertwine intricately, playing important roles in maintaining heart function. This inherent characteristic makes the heart particularly susceptible to iron imbalance, which may significantly contribute to the development of various cardiovascular diseases (Thygesen et al., 2019). Recent investigations have underscored the role of iron imbalance as a recognized factor in MI (Griffiths et al., 1985; Lakhal-Littleton et al., 2015, 2016, 2017, 2019; Weidmann et al., 2020; Dharmakumar et al., 2021; Kalisch-Smith et al., 2021; Jayakumar et al., 2022). As altered iron homeostasis leads to increased reactive oxygen species (ROS) generation via the Fenton reaction, sustaining inflammatory conditions that contribute to MI (Liu et al., 2020; Ying et al., 2022). Therefore, it is utmost crucial to regulate the iron availability to the heart. Recent investigations have explored the understanding of how HEP/FPN operates differently in systemic and cardiac microenvironments. Locally synthesized hepcidin within the heart acts as an autocrine factor, tightly controlling iron levels. Notably, in response to hypoxia, systemic hepcidin levels decrease while cardiac hepcidin levels rise, thereby preserving cellular iron (Lakhal-Littleton et al., 2016). This heightened cardiac hepcidin expression inhibits ferroportin-1 (FPN-1), the sole protein responsible for iron export, effectively retaining iron within the heart. Moreover, targeted deletion of FPN-1 in cardiac cells leads to iron accumulation exclusively within cardiomyocytes, culminating in cardiac dysfunction (Lakhal-Littleton et al., 2015). Hence, elucidating the upstream signaling mechanisms governing the intricate functions of cardiac hepcidin and FPN-1 is of paramount importance. NF-E2-related factor 2 (Nrf2), a member of the basic leucine zipper (b-Zip) transcription factor family, holds a central role in shielding cells from oxidative stress by upregulating gene transcription through the antioxidant-responsive element (ARE). Beyond its antioxidant regulatory function, Nrf2 also exerts significant control over key genes and proteins involved in iron metabolism, hypoxia, and autophagy across various pathophysiological conditions, including MI (Erkens et al., 2015; Qin et al., 2016; Kasai et al., 2018; Liu et al., 2020; Tian et al., 2022). A study by Zhu et al.2008 provided initial evidence of Nrf2-dependent cytoprotection against oxidative and electrophilic stress in cardiomyocytes, utilizing neonatal mouse ventricular myocytes from Nrf2 knockout mice. Mechanistically, Nrf2 activation may bolster antioxidant defenses, regulate metabolism, and modulate autophagy and proteasome function, thus contributing to cardiac protection (Qin et al., 2016). However, contrary to the beneficial effects attributed to Nrf2, Allwood et al. (2014) discovered that transgenic overexpression of HMOX1, a well-established downstream target of Nrf2 in the heart, leads to the spontaneous development of heart failure and exacerbates pressure overload-induced cardiomyopathy in mice. Consequently, Nrf2 exhibits both protective and detrimental effects in cardiovascular diseases. Despite these findings, the precise molecular mechanism underpinning the dual role of Nrf2 signaling in MI remains elusive.

Given Nrf2’s pivotal role in regulating iron metabolism, we hypothesized that myocardial infarction pathogenesis perpetuates Nrf2 activation, thereby disrupting iron homeostasis *via* the hepcidin/ferroportin axis. This disruption instigates ferritinophagy and subsequent intracellular iron accumulation, culminating in cardiomyocyte death. To test this hypothesis, we employed both *in vivo* and *in vitro* approaches. In the *in vivo* model, MI was induced using isoproterenol (ISO), while in the in vitro experiment, differentiated H9c2 cardiomyocytes were subjected to cobalt chloride (CoCl_2_)-induced hypoxic stress, with Nrf2 pharmacological suppression using brusatol. Our findings revealed that enhanced activation of the Nrf2 signalling pathway serves as a crucial mediator in the pathogenesis of MI.

## 2. METHODS

### 2.1. Chemicals and reagents

The chemicals, drugs, antibodies, commercial assay kits, and nucleic acid purification kits used in the current study are listed in the Key Resource Table. All chemicals used were of analytical grade, unless otherwise stated.

### 2.2. *In vivo* studies

Age-matched (8 weeks old) Wistar male and female albino rats, weighing approximately 200 ± 30 g, were procured from the Central Animal House Facility, University of Madras, Taramani Campus, Chennai (Tamil Nadu). Animal experiments were carried out according to the guidelines of the Committee for the Purpose of Control and Supervision of Experiments on Animals (CPCSEA), India, and approved by the Institutional Animal Ethics Committee (IAEC), University of Madras, Tamil Nadu **(Approval No. 01/08/2020 & 01/17/2023).** The myocardial infarction model was established by subcutaneous injection of isoproterenol hydrochloride (ISO) dissolved in sterile water at a dose of 85 mg/kg body weight. For dose-dependent studies, we used 45, 85, and 170 mg/kg of body weight. Induction was carried out for two consecutive days with a 24-hour interval (1, 2). At the end of the experimental period, the rats were anesthetized with ketamine (22 mg/kg body weight, i. p.). Hearts were excised, washed with ice-cold physiological saline, and stored at -80°C for biochemical and molecular studies.

#### 2.2.1. Histopathology and immunohistochemistry

For histopathological studies, the animals were anesthetized with ketamine (22 mg i. p.), the heart was removed and fixed in 4% PFA and allowed to impregnate in 30% sucrose solution in PBS. Serial sections of hearts (apex to base) were cut on a microtome (30µm thickness) and collected on gelatin-subbed slides using PBS.

##### 2.2.1.1. Masson’s trichrome staining

To determine the extent of fibrosis in the heart, paraffin-embedded sections were stained with Masson’s trichrome stain. Briefly, heart sections from both the control and ISO-induced MI animals were stained with Bouin’s solution for 30 min, stained with Weigert’s hematoxylin for 20 min, and incubated with Biebrich scarlet-acid fuchsin solution for 20 min. Tissue sections were then treated with phosphomolybdic-phosphotungstic acid solution for 5 min, followed by aniline blue solution for 2 min. After incubation with 1% acetic acid for 1 min, sections were dehydrated with graded alcohol and xylene. Fibrotic areas (blue-stained) within the sections were visualized using a light microscope (Accu-scope 2019 EXC-500 Upright Microscope).

##### 2.2.1.2. Perl’s Prussian blue staining

To assess the deposition of nonheme iron, such as ferritin and hemosiderin, in both control and ISO-group, heart sections were treated with 2% hydrochloric acid, followed by 2% potassium ferrocyanide, and incubated for 30 min. The tissue sections were then rinsed thrice with distilled water for 1-2 minutes. The sections were dehydrated using graded alcohol and xylene solutions, and the slides were washed with 0.1M PBS. Sections were counterstained with 3,3-diaminobenzidine for 5 min at room temperature. The iron deposited in the tissue may subsequently be visualized as a complex hydrated ferric ferrocyanide substance that appears as an insoluble purple/Prussian blue pigment under a light microscope (Accu-scope 2019 EXC-500 Upright Microscope).

##### 2.2.1.3. Immunohistochemistry

Myocardial sections from control and ISO-induced MI rats were deparaffinized, washed with PBS, and incubated with 10% goat serum in PBS for 2 h. After blocking, sections were incubated with primary antibodies against Nrf2 and hepcidin (rabbit monoclonal, 1:250) in 10% goat serum containing 0.2% Triton X-100 for 24 h at 4°C on a platform shaker. After rinsing in PBS, the sections were incubated with secondary anti-rabbit IgG-conjugated horseradish peroxidase antibody (1:1000) in PBS (pH 7.4) for 60 min at room temperature. Visualization was performed by incubation in 3,3-diaminobenzidine (DAB) for 5 min, after which the sections were examined under a light microscope (Accu-scope 2019 EXC-500 Upright Microscope).

#### 2.2.2. Analysis of cytokines using enzyme linked immunosorbent assay (ELISA)

The levels of serum inflammatory cytokines (CRP, TNF-α, IL-6, IL-8, and IL-1α) in the control and ISO-induced MI animals were estimated using ELISA. Briefly, 50 μg of protein from the serum sample was coated onto 96 well plates using bicarbonate buffer and left overnight at 4°C. The next day, plates were washed with PBS and blocked with 1% BSA solution (200 μl/well) for 2 h to prevent non-specific binding in the subsequent steps. The wells were then washed four times with PBS and dried. The specific primary antibody, diluted in 1% BSA (1:1000) was added to the wells (100μl/well) and incubated overnight at 4°C. The following day, the wells were washed with PBS-Tween 20 four times and incubated with the secondary conjugated horseradish peroxidase antibody for an hour at room temperature. After washing, 100 μl of ABTS substrate was added and incubated in the dark for 20 min. Then, 2 N sulfuric acid (100μl/well) was added to the wells to stop the reaction, and the plates were read at 415 nm using a Bio-Rad iMark™ Microplate Absorbance Reader (Bio-Rad Laboratories, Hercules, CA, USA).

#### 2.2.3. Iron estimation from myocardial tissue

Briefly, frozen heart tissue was cut into pieces and dried overnight at 106°C in a 35mM dish. The dried tissue pieces were weighed, immersed in a 0.5 ml acidic mixture (containing 3 M HCl and 10% TCA), and incubated at 65°C for 42 h. The obtained lysates were centrifuged at 10,000×g for 10 min, and the supernatants were collected in different tubes. From this, 50 µL of each sample was placed on a 96-well plate, and 200-μl iron-detection reagent (6.5mM Ferrozine; 6.5mM Neocuproine; 2.5M ammonium acetate; 1M ascorbic acid) was added to each well. Following a 30-min incubation at room temperature, the absorbance was recorded at 562 nm. The amount of iron per milligram of dry tissue was calculated from a standard curve using FeCl_3_ (Daba et al 2013).

#### 2.2.4. Determination of myocardial glutathione redox state

The level of reduced glutathione (GSH) was determined using the method described by Moron et al. (1979) with slight modifications. In brief, myocardial lysates were prepared in 2-(N-morpholino) ethanesulfonic acid (MES) buffer and centrifuged at 5000 rpm for 5 min at 4°C. A small aliquot of the supernatants was used for protein determination and the remaining samples were mixed with equal volumes of 10% meta-phosphoric acid to precipitate proteins. A known volume of the metaphosphoric acid extracts was treated with triethanolamine reagent and 2-vinyl pyridine (only for GSSG analysis) and appropriate GSH and GSSG standards were treated similarly to prepare a standard graph. This method is based on the reaction of GSH with DTNB to form TNB, which absorbs light at a wavelength of 412 nm. The level of glutathione was expressed as nmol/mg protein.

#### 2.2.5. Analysis of lipid peroxidation and protein carbonyls in heart tissue

Lipid peroxide levels were measured following the method of Devasagayam and Tarachand (1987). The results were expressed as µmoles of MDA per milligram of protein. Protein carbonyl levels in the tissue were assessed using 2,4-dinitrophenyl hydrazine (DNPH), as described by Levine et al. (1990), with slight modifications. The absorbance difference between DNPH-treated and HCl-treated samples was measured at 366 nm. An extinction coefficient of 22.0 mM⁻¹cm⁻¹ was used to calculate the values, which were expressed as µmoles of protein carbonyl per milligram of protein.

#### 2.2.6. RNA isolation and qPCR

Total RNA was isolated from myocardial tissues using TRIzol reagent and a RNeasy mini kit according to the manufacturer’s instructions. The purity of the RNA samples was assessed and quantified by measuring the absorbance ratio (A) at 260/280 using a Nano spectrophotometer (Thermo Fisher Scientific Inc., United States). Total RNA (2 μg) was subjected to cDNA synthesis using the iScript cDNA Synthesis Kit (Bio-Rad Laboratories, Hercules, CA, USA). Real-time PCR was performed on an Applied Biosystems 7300 Sequence Detection System (Applied Biosystems, Foster City, CA, USA) using the Powerup™ SYBR Green Master Mix (Thermo Fisher Scientific Inc., United States). The primer sequences used for gene amplification were procured from Sigma– Aldrich (St.Louis, MO, USA). Louis, MO, USA) and Eurofins Genomics (Ebersberg, Germany), and are provided in the Key Resources Table. Gene expression levels were normalized to those of β-actin. The fold change in gene expression indicated by all PCR results was calculated using the 2^−ΔΔCt^ method.

### 2.3. *In Vitro* studies using H9c2 cells

H9c2 (2-1) BDIX Heart Myoblast cells were obtained from the National Center for Cell Science, Pune, India. The cells were initially grown in DMEM supplemented with 10% fetal bovine serum (*v*/*v*),100 U/ml penicillin, and 100 μg/ml streptomycin in 25 cm^2^ and 75 cm^2^ vented culture flasks. Cultures were incubated at 37°C in 5% CO2/95% humidified air. When the cells reached 80–90% confluence in the flask, they were trypsinized and seeded onto 96-well or 6-well plates. To induce differentiation, H9c2 cells were transferred to a differentiation medium (DMEM containing 0.5% FBS, Pen/Strep) with 2.5 μM all-trans retinoic acid (ATRA) for 5 days (Lin et al., 2010). The differentiation media (DM) was replenished and fresh ATRA was added every alternative day. After the treatment period, the differentiated cells were transferred to a proliferation medium (PM) in 6-well or 60 mm dishes and used for various experiments.

#### 2.3.1. Immunofluorescence, image acquisition, and quantification

H9c2 cells were cultured on coverslips pre-coated with 0.02% gelatin (in 6-well plates) and treated with 2.5 μM all-trans retinoic acid (ATRA) for 5days. For staining, the cells were incubated with 4% paraformaldehyde for 15 min, washed thrice with PBS, and permeabilized with 0.25% Triton X-100. After three washes with PBS, the cells were blocked with 5% goat serum for 1 h and then incubated with cardiac troponin-T antibody (1:250, mouse monoclonal in 5% goat serum) in a light-protected chamber maintained overnight at 4°C. Following incubation, the cells were washed thrice with PBS and incubated with Alexa fluor 594 conjugated secondary antibody for 1 h at RT. The cells were then washed three times with PBS and mounted with fluoroshield/DAPI. Images were acquired with an Accu-scope 2019 EXC-500 fluorescence microscope using a 20/40× objective lens.

#### 2.3.2. Cell viability using MTT assay

To determine the optimal dose of cobalt chloride (CoCl_2_) and brusatol, cell viability was evaluated using the 3-(4,5-dimethylthiazol-2-yl)-2,5-diphenyl-tetrazolium bromide (MTT) assay (Kumar et al., 2018). Differentiated H9c2 cardiomyocytes (5 x 10^3^ per well) were plated in 96-well flat-bottomed culture plates. The cultures were treated with CoCl_2_ (10 – 500 μM) or brusatol (10 – 100 nM) for 24 h. After treatment, MTT reagent was added to each well, and the plate was incubated for 4 h at 37°C. Dimethyl Sulfoxide (DMSO) was added and the optical density (OD) of the sample plate was measured at 510 nm in a microplate reader. The results for each experimental condition were verified a minimum of three times. Based on cell viability, the optimum concentration of CoCl_2_ was found to be 100 μM, while the optimal dosage for brusatol was found to be 50nM. For *in vitro* experiments, differentiated H9c2 cells were pretreated with brusatol (50nM) for 3 h and/or CoCl_2_ (100μM) for 24 h.

#### 2.3.3. Iron estimation from H9c2 cardiomyocytes

After the treatment period, H9c2 cardiomyocytes were washed thrice with PBS. 200μL of 50mM NaOH was added to each well, and the plates were incubated at -20°C overnight. The next day, the cells were lysed, and aliquots were collected in fresh 1.5mL tube. To 100μL of lysate, 100μL of 10mM Hcl and 100μL of iron-releasing agent (4.5% KMnO4 in 1.4M Hcl) were added and incubated at 60°C for 2-hours. Subsequently, the samples were cooled to room temperature. To this, 30μL of iron-detection reagent (6.5mM Ferrozine + 6.5mM Neocuproine + 2.5M ammonium acetate + 1M ascorbic acid) was added and incubated for 30 mins at RT. 280μL of this reaction mixture was transferred into a 96-well plate and absorbance was recorded at 562 nm in a microplate reader. The iron standards (FeCl_3_) were treated in the same manner. Protein estimation in the lysates was performed separately.

#### 2.3.4. Analysis of apoptotic cells by flow cytometry

The different stages of apoptosis in the control, CoCl_2_ induced hypoxic and CoCl_2_ + brusatol treated H9c2 cells were analyzed by flow cytometry using an Annexin V-FITC/PI Apoptosis kit (ElabScience, Houston, USA). After the incubation period, cells were washed with cold phosphate-buffered saline (PBS), centrifuged twice at 1500 rpm for 5 min, and suspended in 100μL of binding buffer. FITC-labeled annexin V (2.5μL) and propidium iodide (PI, 2.5μL) were added and incubated with the cells at room temperature for 20 min. Apoptotic cells were measured using a FACSCalibur flow cytometer (Beckman Coulter Cytoflex, California, USA). Annexin V-positive and PI-negative cells were scored as early apoptotic. Cells that were positive for both annexin V and PI were considered to be late apoptotic cells.

### 2.4. Immunoblotting

Heart tissue lysates and H9c2 cell lysates (50 μg protein) were separated by SDS-PAGE on 10– 12% polyacrylamide gels and transferred to polyvinylidene difluoride (PVDF) membranes. The membranes were incubated with specific primary antibodies (listed in the key resources table) for approximately 16 hours (overnight) at 4°C. After washing thrice with TBST, the membranes were incubated with horseradish peroxidase-conjugated secondary antibodies for 1 h. Immunoreactive bands were developed using Clarity™ Western ECL Substrate (with an equal volume of luminol and H_2_O_2_) and visualized using a ChemiDoc XRS imaging system (Bio-Rad Laboratories, CA, USA). To verify the uniformity of the protein load and transfer efficiency across the test samples, the membranes were reprobed with β-actin and Lamin B.

### 2.5. Data and statistical analyses

All experiments were designed to generate groups of equal size using randomization and blinded analyses. Group size was the number of independent values, and statistical analyses were performed using these independent values. All datasets were tested for normal distribution and homogeneity of variance to confirm that nonparametric testing was not required. Statistical analyses were performed using Prism 8.0 (GraphPad Software, San Diego, CA, USA). Data are shown as mean ± SEM. A comparison between the two groups was performed using the unpaired t-test. Multiple group comparisons were performed using ANOVA followed by the Bonferroni procedure for the comparison of means. Differences among means were considered statistically significant when the p-value was less than 0.05.

## 3. RESULTS

### 3.1. Establishing isoproterenol-induced myocardial infarction in Wistar rats

Isoproterenol (ISO), a synthetic catecholamine and β-adrenergic agonist, is administered in varying doses to rodents and mimics diverse human cardiovascular diseases. In our previous study, we established an ISO-induced cardiac hypertrophy model using a dose of 5 mg/kg body weight (Velusamy et al., 2020). Various other investigations have shown that ISO (85 and 100 mg/kg body weight), when subcutaneously injected into rats at 24-hour intervals for two days, exhibits myocardial infarction (MI) (Ekici et al., 2022). Based on this, we conducted dose-dependent studies to identify the optimal ISO concentration for mimicking myocardial infarction (MI) by administering doses of 45, 85, and 170 mg/kg subcutaneously to adult Wistar rats over two days (Figure S1). Cardiac histopathological analysis of control rats revealed regular arrangements of myofibrils with clear striation, while the 45mg dose exhibited wavy arrangement of myofibril with mild infiltration along with undistributed nuclei pattern. Both the 85 and 170 mg displayed disoriented myofibrils with neutrophil infiltration and necrotic impressions, however extensive subendocardial necrosis along with intramyofiber edema and vacuoles were also found enormous in the later (Figure S1A). Levels of Creatine kinase (CK) and lactate dehydrogenase (LDH), were assessed to evaluate the cardiac tissue degeneration. Additionally, changes in systemic iron metabolism were evaluating the serum iron and hepcidin levels systemic iron regulation in all the dosages. No significant changes were observed in 45 mg, on the other hand both the 85 and 170 mg dosages caused a significant elevation in all the parameters with subsequent decreased levels of serum iron in all the groups when compared to control (Figure S1B S1C). Based on these evidences, 85mg is considered to an optimum dose to induced MI due to its ability to mimic the early stages of MI with less evidence of necrosis and lower animal mortality rates.

Figure 1A outlines the study design. Gross examination of the heart revealed a considerable increase in cardiac mass in ISO-treated rats (Figure 1B). Similarly, four-chambered heart images showed a robust increase in the cardiac mass, in particular left ventricle showed a pathological remodeling (Figure 1C). Further. morphological examination exhibited higher heart weight to body weight (HW/BW), left ventricle to body weight (LV/BW), and LV/tibia length ratios in ISO-induced rats when compared to control rats (Figure 1D). The serum levels of cardiac troponin T (cTnT) was measured using ELISA. The results showed a significant increase, approximately 2-fold, in the serum of MI-induced rats compared to the control (Figure 1E). Histopathological analysis of cardiac tissue from control rats demonstrated regular arrangements of myofibrils with distinct striations, while the rats administered with ISO 45 mg revealed wavy arrangements of myofibrils with mild neutrophil infiltration and undistributed nuclei pattern. However, disoriented myofibrils with damaged striations, intense neutrophil infiltration and nuclei aggregation were observed in both 85 and 170 mg doses (Figure 1F). Masson’s trichrome staining revealed elevated collagen deposition in ISO-induced animals, coupled with disoriented myofibrils, neutrophil infiltration, and necrotic impressions, confirming fibrotic degeneration of the cardiac tissue (Figure 1G and S1). To determine the extent of neutrophil infiltration and inflammatory changes caused by ISO, levels of inflammatory cytokines in serum were measured using ELISA (Figure 1H). The levels of CRP, TNF-α, IL-6, IL-8, and IL-1β significantly increased (P < 0.001) following ISO treatment, confirming the induction of MI in these rats. Overall, histopathological, and biochemical analyses confirmed the establishment of MI through the subcutaneous induction of ISO.

**Figure 1:**
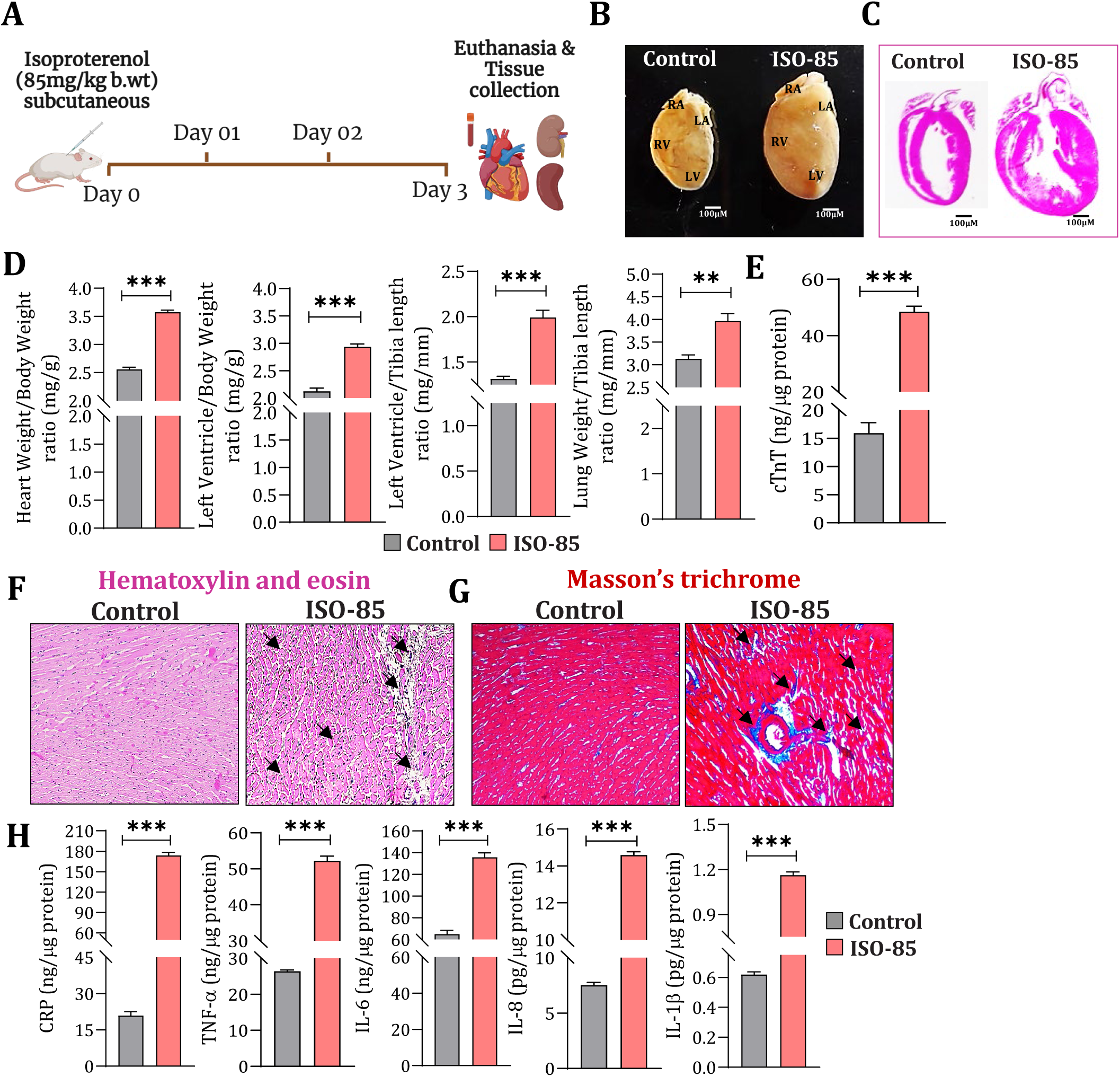
Establishing isoproterenol-induced myocardial infarction in Wistar rats. **(A)** Schematic representation of the study design. **(B)** Gross examination of the heart from Control and ISO-induced rats revealed that cardiac mass had increased considerably in ISO-treated rats. **(C)** Representative imageof four chambered sections of the myocardium from control and ISO-administered rats. **(D)** Bar graph depicting the ratio of heart weight/body weight (HW/BW, mean 2.560 vs. 3.575, p < 0.001), left ventricle/body weight (LV/BW, mean 2.12 vs. 2.93, p < 0.001), left ventricle/tibia length (LV/TL, mean 1.314 vs. 1.993, p < 0.001), and lung weight/tibia length (LW/TL, mean 3.13 vs. 3.96, p = 0.0021), n = 6 per group. **(E)** Bar graph depicting the cardiac troponin T (cTnT) levels in the serum of control and ISO-induced myocardium measured by ELISA (mean 15.91 vs 48.41, p < 0.001), n = 6 per group. **(F)** Paraffin sections showing inflammatory infiltration (black arrows) in the ISO myocardium compared to the control by H&E staining (n = 5 per group). **(G)** Paraffin sections showing fibrotic degeneration and enhanced collagen deposition in the ISO myocardium (black arrows) compared to the control by Masson Trichrome staining (n = 5 per group). **(H)** Bar graphs depicting the levels of inflammatory cytokines in the serum of control and ISO-induced myocardium measured through ELISA: CRP (mean 20.88 vs 173.8, p < 0.001), TNF-α (mean 26.42 vs 52.24, p < 0.001), IL-6 (mean 64.99 vs 135.70, p < 0.001), IL-8 (mean 7.52 vs 14.59, p < 0.001) and IL-1β (mean 0.61 vs 1.16, p < 0.001), n = 6 per group. All data are presented as the mean ± SEM. Statistical significance was calculated using an unpaired t-test with Welch’s correction (*p < 0.05; **p < 0.01; ***p < 0.001).

### 3.2. Isoproterenol-induced MI leads to alterations in cardiac iron metabolism

Recent studies have drawn attention to the importance of dysregulated iron metabolism in cardiac patients (Ghafourian et al., 2020; Lavoie, 2020; Ito et al., 2021). Investigating dynamic changes in iron homeostasis following ISO-induced MI and their role in the disease’s progression is therefore crucial. The serum iron levels were found to be significantly decreased (p < 0.001) while hemoglobin levels remained unaffected. Conversely, hepcidin and ferritin levels were substantial increase (p < 0.001) in ISO-induced MI rats (Figure 2A,2B). To delve deeper into the transient changes in the key protein of iron metabolism in myocardium of ISO induced animals, qPCR and immunoblot analyses were performed. Initially, the gene expression and protein level of hypoxic inducible factor were studies. Both Hif1α & Hif2α gene levels were augmented (p=0.002 & 0.0033) respectively. In line with this, the gene expression of *Epo*, a hypoxia-responsive gene, was also elevated upon ISO-induced MI (Figure 2C). While the immunoblot, analysis revealed elevated nuclear HIF-1α levels with declined cytosolic HIF-1α upon ISO induced, suggested a potential hypoxic environment compared to control rats (P < 0.05) (Figure 2E-F). Next the gene expression of iron metabolism proteins; transferrin (Tf) receptor (TfR), iron regulatory proteins (Hamp, Slc40a1/HEP/FPN-1) and its regulatory targets (Bmp6, HJV, Stat3, Smad1 and Smad4), Epo a hormone responsible for RBC maintenance and finally iron storage protein (FTL and FTH) was assessed. Intriguingly, the levels all of these transcripts were found to be augmented except Smad4 with maximum upregulation in *Slc40a1* (∼7-fold) and *Smad1* (∼8-fold) in ISO induced rats (Figure 2C). However, at the protein level no discernible differences in FPN-1 and a notable two-fold upregulation of hepcidin were observed in the ISO myocardium, as confirmed by western blotting (Figure 2E-F) and immunohistochemistry (Figure 2D), in comparison to the control. Interestingly, while both *Fth* and *Ftl* gene expression levels increased significantly, FTH protein levels were found to be downregulated (P < 0.001), whereas FTL levels (Figure 2E-F) were elevated in myocardial lysates (P < 0.01). Overall, these findings provide conclusive evidence of alterations in iron metabolism during ISO-induced MI, highlighting specific modifications in FTH, FTL, hepcidin, and FPN-1.

**Figure 2:**
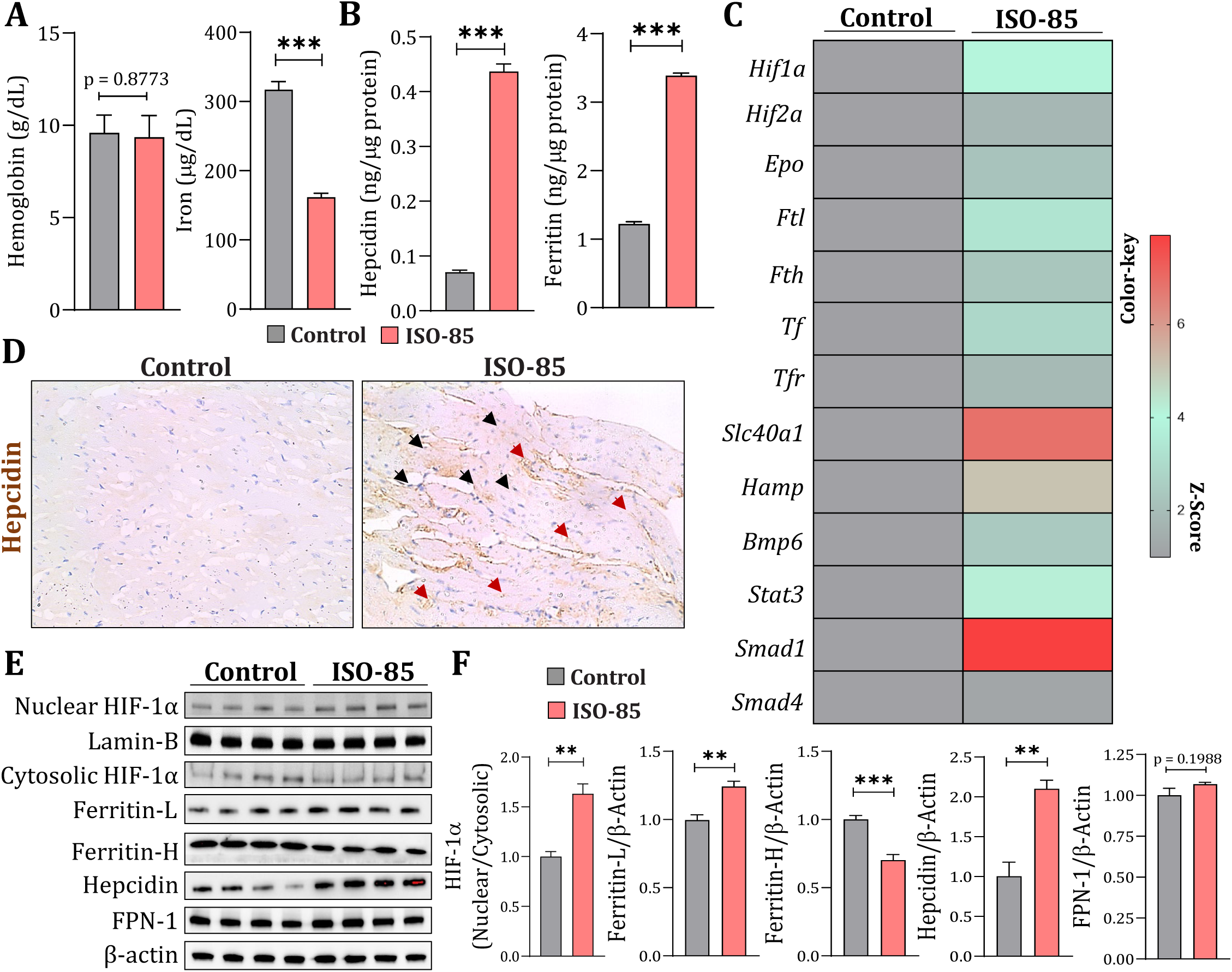
Isoproterenol-induced MI leads to alterations in cardiac iron metabolism. **(A)** Bar graphs depicting hemoglobin and iron levels in the serum of control and ISO-induced myocardium. No significant changes were observed in hemoglobin levels (mean 9.596 vs. 9.352, p = 0.8773); however, iron levels were significantly downregulated in ISO-induced MI compared to the control (mean 317.2 vs 161.8, p < 0.001), n = 5 per group. **(B)** Bar graphs depicting hepcidin and ferritin levels in the serum of the control and ISO-induced myocardium. A significant increase in hepcidin (mean 0.070 vs. 0.43, p < 0.001) and ferritin (mean 1.22 vs. 3.38, p < 0.001) levels was observed in the ISO-induced group when compared to the control group (n = 6 per group). **(C)** qPCR of hypoxia/iron metabolism/Stat3/Bmp6 signaling genes in the control and ISO-induced myocardium. Fold changes (relative quantification, RQ) were normalized to the endogenous *Actb* (β-actin) expression. Differential expressions of hypoxia [(*Hif1a,* mean 1.0 vs 4,50, p = 0.0006), (*Hif2a,* mean 1.0 vs 1.89, p = 0.0034) and (*Epo,* mean 1.0 vs 2.35, p < 0.001)], iron metabolism [(*Ftl,* mean 1.0 vs 3.63, p < 0.001), (*Fth,* mean 1.0 vs 2.45, p < 0.001), (*Slc40a1,* mean 1.0 vs 6.99, p < 0.001), (*Tf,* mean 1.0 vs 3.13, p = 0.0064), (*Tfr,* mean 1.0 vs 1.98, p = 0.0004), and (*Hamp,* mean 1.0 vs 5.46, p = 0.0013)] and Stat3/Bmp6 [(*Bmp6,* mean 1.0 vs 2.59, p = 0.0019), (*Stat3,* mean 1.0 vs 4.61, p = 0.0006), (*Smad1,* mean 1.0 vs 7.91, p < 0.001), and (*Smad4,* mean 1.0 vs 1.20, p = 0.1321) signaling genes is illustrated as a heat map (n = 6 per group). **(D)** Immunohistochemical staining of hepcidin protein in control and ISO-induced rat heart samples revealed enhanced localization of hepcidin in the ISO-myocardium compared to the control, as indicated by the black arrows (n = 5 per group). **(E)** Immunoblot analysis of nuclear and cytosolic HIF1α, Ferritin-L, Ferritin-H, Hepcidin and FPN-1 in control and ISO-induced myocardial lysates. ISO-induced MI leads to elevation in HIF1α, Ferritin-L, and Hepcidin levels. In addition, ferritin-H was downregulated in ISO-induced animals, and no significant difference was found in FPN-1 protein levels between control and ISO-induced animals. **(F)** Respective densitometries of HIF1α (Nuclear/cytosolic ratio, mean 1.0 vs 1.63, p = 0.0015), Ferritin-L (mean 1.0 vs 1.24, p = 0.0019), Ferritin-H (mean 1.0 vs 0.70, p = 0.0006), Hepcidin (mean 1.0 vs 2.10, p = 0.0014), and FPN-1 (mean 1.0 vs 1.068, p = 0.1988), n = 5 per group. All data are presented as the mean ± SEM. Statistical significance was calculated using an unpaired t-test with Welch’s correction (*p < 0.05; **p < 0.01; ***p < 0.001).

### 3.3. Alterations in cardiac iron metabolism during MI are due to augmented Nrf2 transcriptional activity

Nuclear factor erythroid 2-related factor 2 (Nrf2) and antioxidant response element (ARE) (Nrf2/ARE) signalling pathway is reported as a prime transcriptional regulator of most of the key proteins involved in iron metabolism (Bayele et al., 2015; Kerins & Ooi, 2018; Saha et al., 2020). Building on this, we comprehensively examined the changes in Nrf2 and its downstream targets both at the transcriptional and translational level. The gene expression of Nrf2 (*Nfe2l2*) observed a four-fold increase (Figure 3A) while, immunostaining observation exhibited localization of Nrf2 in the nucleus was more in ISO-induced MI animals than the control (Figure 3B). These findings were further supported by immunoblotting, which demonstrated that elevated nuclear accumulation of Nrf2 along with a significant decrease in cytosolic levels (nuclear/cytosolic ratio: mean 1.0 vs 4.72, p <0.001) (Figure 3C, 3D). This finding emphasizes the translocation of Nrf2 to the nucleus in response to ISO.

**Figure 3:**
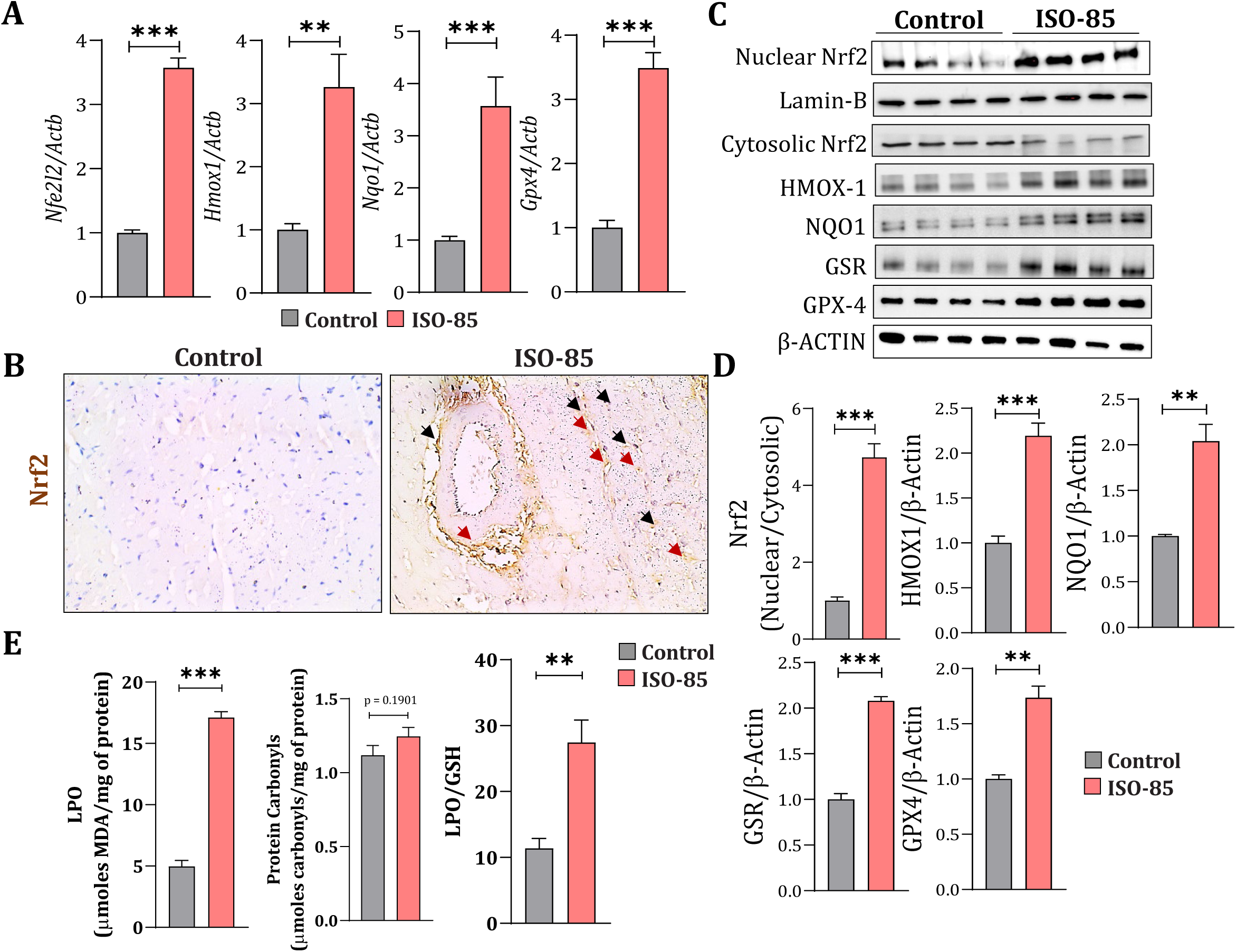
Alterations in cardiac iron metabolism during MI are due to augmented Nrf2 transcriptional activity. **(A)** qPCR of Nrf2 and its downstream targets in control and ISO-induced myocardium. Fold changes (relative quantification, RQ) were normalized to the endogenous *Actb* (β-actin) expression. Differential expression of Nrf2 (*Nfe2l2,* mean 1.0 vs 3.57, p < 0.001), *Hmox1* (mean 1.0 vs 3.26, p = 0.0068), *Nqo1* (mean 1.0 vs 3.56, p = 0.0056), and *Gpx4* (mean 1.0 vs 3.48, p < 0.001) are illustrated as bar graphs (n = 6 per group). **(B)** Immunohistochemical staining of Nrf2 in control and ISO-induced rat heart samples revealed enhanced nuclear localization of Nrf2 in the ISO-myocardium compared to the control, as indicated by the black arrows (n = 5 per group). **(C)** Immunoblot analysis of nuclear and cytosolic Nrf2, HMOX-1, NQO1, GSR, and GPX4 in control and ISO-induced myocardial lysates. ISO-induced MI leads to an elevation in the nuclear levels of Nrf2, whereas the cytosolic levels of Nrf2 decrease. Similarly, the downstream targets HMOX-1, NQO1, GSR, and GPX4 were enhanced in ISO-induced myocardium. **(D)** Respective densitometries of Nrf2 (nuclear/cytosolic ratio, mean 1.0 vs 4.72, p = 0.0003), HMOX-1 (mean 1.0 vs 2.19, p = 0.0003), NQO1 (mean 1.0 vs 2.04, p = 0.0046), GSR (mean 1.0 vs 2.08, p < 0.001), and GPX4 (mean 1.0 vs 1.736, p = 0.0012) (n = 5 per group). **(E)** Bar graph depicting lipid peroxidation (LPO), protein carbonyls and ratio of lipid peroxidation to reduced glutathione (LPO/GSH) in the myocardium of control and ISO-administered animals. A significant increase in LPO levels (p < 0.001) was observed. No significant change in the protein carbonyl levels (p = 0.1901) was observed. A significant increase (p = 0.0036) in the ratio (LPO/GSH) was observed in the ISO administered animals compared to control, n = 6 per group. All data are presented as the mean ± SEM. Statistical significance was calculated using an unpaired t-test with Welch’s correction (*p < 0.05; **p < 0.01; ***p < 0.001).

To corroborate the enhanced Nrf2 transcriptional activity, further investigation included the evaluation of downstream Nrf2 targets at the gene and protein levels. Our findings demonstrated elevated mRNA and protein levels of HMOX1, NQO1, GPX4 and protein levels of GSR in ISO-induced rats (P < 0.05), indicating enhanced Nrf2 transcriptional activity (Figure 3A, 3C-D). Expanding on our investigation, we assessed the levels of reduced glutathione (GSH), lipid peroxidation (LPO), and protein carbonyls (PC) in control and ISO-administered rats. Interestingly, there was a ∼3.5-fold upregulation in LPO levels, while no significant changes (p = 0.1901) were observed in protein carbonyl levels in the ISO-administered myocardium. Further, ratio of malondialdehyde (MDA or LPO) to GSH revealed a significant increase in ISO-induced MI animals (Figure 3E) when compared to control animals denoting that the myocardium is in the oxidative state despite active antioxidant abundance. These results indicate that elevated Nrf2 levels contribute to redox alterations in the myocardium, primarily aligning with myocardial damage and degeneration induced by ISO.

### 3.4. Augmented Nrf2 transcriptional activity leads to increased iron sequestration and subsequent ferritinophagy

Emerging evidence have shed light on the mutual crossover of Nrf2 and iron metabolism regulation in the progression of myocardial infarction (Jayakumar et al., 2022); therefore, we further investigated the downstream consequences of augmented Nrf2 on altered iron metabolism in ISO induced MI rats. Increased activation of HMOX-1 following ISO treatment is anticipated to increase heme degradation, thereby contributing to elevation of the labile iron pool (LIP). Concurrently, the colorimetric ferrozine assay and Perl’s Prussian blue staining were performed to examine the intracellular iron content levels in the myocardium of experimental rats. Consistent with the augmented hepcidin levels, our observations revealed a substantial increase in iron accumulation, manifested by numerous blue/purple granular spots, demonstrating iron-containing hemosiderin/ferritin molecules and the release of Fe^3+^ ions (black arrows) in rats subjected to ISO, as opposed to control animals (Figure 4A). Further colorimetric assay also confirmed this elevated iron (P < 0.05) within the myocardium upon ISO-induced MI (Figure 4B). This unequivocally underscores enhanced sequestration of iron in the myocardium. Apparently, Nrf2 is an essential regulator of nonheme-associated iron homeostasis in addition to heme metabolism which include FTL and FTH1 (Cheng H et al., 2020). It is likely that augmented gene expression of both Fth and Ftl (P < 0.05) levels (Figure 2C) can be attributed to increased Nrf2 activity upon ISO induction. Nonetheless, the underlying cause of the downregulation of FTH at the protein level remains undetermined. Conversely, existing literature suggests that FTL and FTH are subject to variable post-transcriptional regulation under different circumstances, necessitating further investigation in ISO-induced rats.

**Figure 4:**
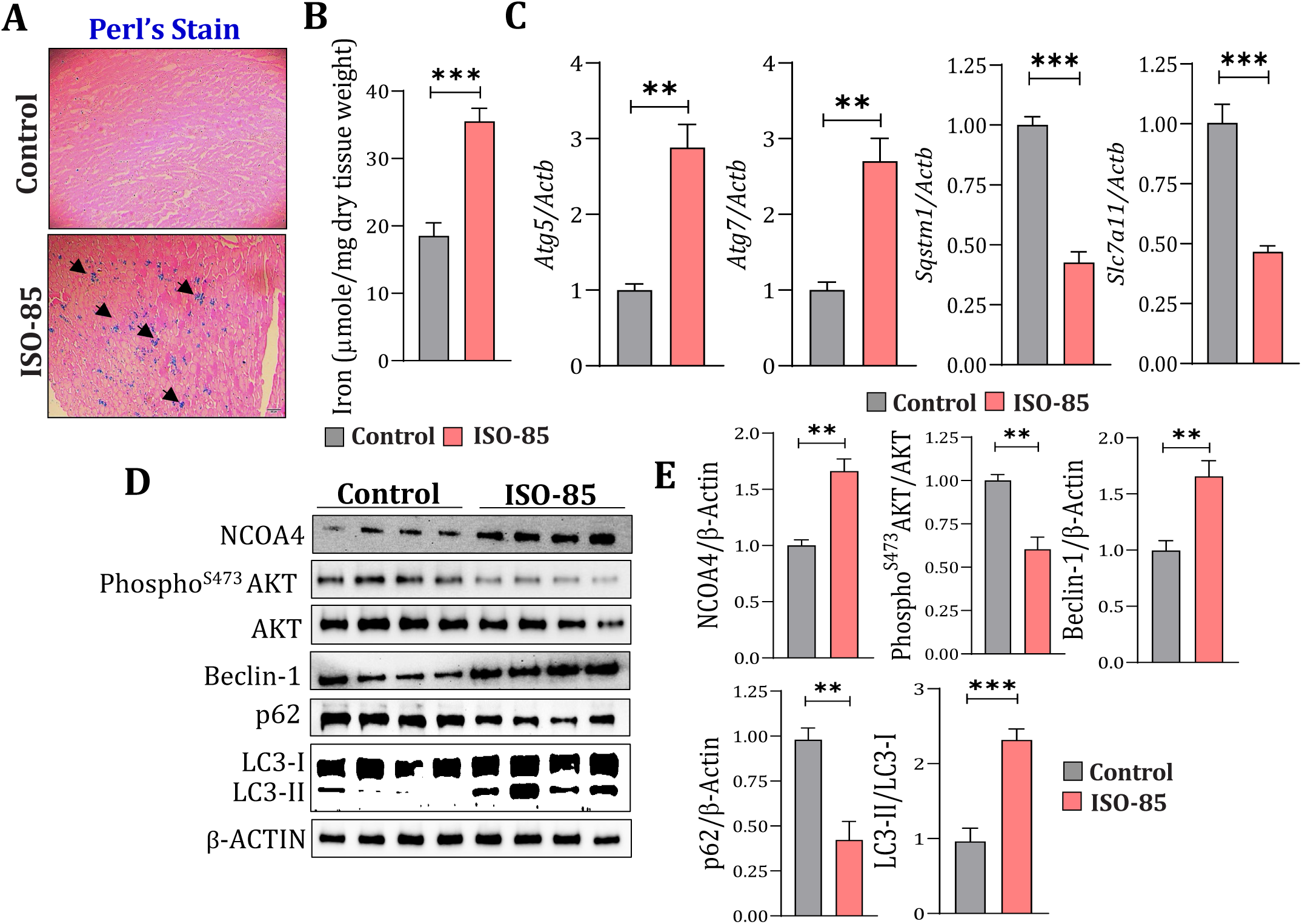
Augmented Nrf2 transcriptional activity leads to increased iron sequestration and ferritinophagy. **(A)** Paraffin sections revealed a substantial increase in iron accumulation, manifested by numerous blue/purple granular spots, demonstrating iron-containing hemosiderin/ferritin molecules and the release of Fe^3+^ ions (black arrows) in rats subjected to ISO, as opposed to control animals (n = 5 per group). **(B)** Bar graph depicting the iron levels measured by the colorimetric ferrozine assay in the myocardium of control and ISO-induced animals. The results matched Perl’s Prussian staining, revealing a significant increase in the levels of iron (mean 18.50 vs 35.50, p < 0.001) within the myocardium of ISO-induced animals when compared to the control (n = 6 per group). **(C)** qPCR of autophagy-related proteins in control and ISO-induced myocardium. Fold changes (relative quantification, RQ) were normalized to the endogenous *Actb* (β-actin) expression. Differential expressions of *Atg5* (mean 1.00 vs 2.87, p = 0.0013), *Atg7* (mean 1.00 vs 2.70, p = 0.0016), *Sqstm* (mean 1.00 vs 0.43, p < 0.001), and *Slc7a11* (mean 1.00 vs 0.47, p < 0.001) illustrated as bar graphs (n = 6 per group). **(D)** Immunoblot analysis of NCOA4, phosphorylated (Serine473) AKT, AKT, Beclin-1, p62, LC3-I, and LC3-II in control and ISO-induced myocardial lysates. ISO-induced MI leads to elevated levels of NCOA4, Beclin-1, and LC3-I/LC3-II, while phosphorylated AKT and P62 were downregulated compared to control animals. **(E)** Respective densitometries of NCOA4 (mean 1.00 vs 1.66, p = 0.0019), AKT (Phospho/Total AKT ratio, mean 1.00 vs. 0.60, p = 0.0023), Beclin-1 (mean 1.00 vs. 1.65, p = 0.0055), P62 (mean 1.00 vs. 0.43, p = 0.0029), and LC3 (LC3-II/LC3-I ratio, mean 1.00 vs. 2.412, p = 0.0004), n = 5 per group. All data are presented as the mean ± SEM. Statistical significance was calculated using an unpaired t-test with Welch’s correction (*p < 0.05; **p < 0.01; ***p < 0.001).

### 3.5. Augmented Nrf2 transcriptional activity leads to increased iron sequestration and subsequent ferritinophagy

To unravel the mechanisms behind increased iron sequestration alongside downregulated FTH levels, we investigated the status of ferritinophagy in rats with ISO-induced myocardial infarction (MI). Ferritinophagy, specialized form of selective autophagic degradation targeting ferritin, facilitated by the nuclear receptor activator 4 (NCOA4) binding to FTH1, thereby regulating intracellular iron bioavailability. Notably, the decline in FTH protein expression (Figure 2E, 2F) coincided with elevated NCOA4 levels (P < 0.05), correlating with increased intracellular iron content upon ISO administration (Figure 4D, 4E). Furthermore, we assessed core protein levels associated with autophagy. Autophagy is regulated by various signaling pathways, including the PI3K/AKT/mTOR pathway (Yu et al., 2015). Thus, a decrease in the ratio of phosphorylated Akt to non-phosphorylated Akt (PhosphoS473Akt/Akt); elevated levels of Beclin-1 and the downstream autophagic effector LC3-II/LC3-I ratio (Figure 4D, 4E) indicated hyperactive autophagy following ISO treatment. Additionally, gene expression of *Atg5* and *Atg7* were elevated (∼3-fold), in contrast *Slc7a11* is decreased in ISO-induced MI when compared to control animals (Figure 4C). Remarkably, the analysis of gene and protein levels of p62 (*Sqstm*) demonstrated a significant (P < 0.01) decrease in ISO-induced MI (Figure 4C-D). Our findings strongly indicate that ISO-induced MI enhances ferritinophagy, supported by elevated levels of NCOA4, other autophagy-related proteins and ferritin degradation.

### 3.5. Pharmacological suppression of Nrf2 rescues hypoxic stress-induced changes in iron metabolism and prevents cardiomyocyte death

To validate the proposed mechanism wherein the altered iron metabolism in the ISO-induced model is primarily mediated by augmented Nrf2-transcriptional activity, we exploited a pharmacological intervention to suppress Nrf2-transcriptional activity in an *in vitro* H9c2 model under hypoxic stress, mimicking MI. Initially, we differentiated H9c2 cardiomyoblasts into cardiomyocytes using All-Trans Retinoic Acid (ATRA, 2.5μM) in a differentiation medium (DMEM containing 0.5% FBS) for 5-days. Differentiation was confirmed by immunostaining and western blotting for cardiac Troponin T (cTnT), a cardiomyocyte marker (Figure 5A, 5B). Subsequently, differentiated H9c2 cardiomyocytes were subjected to hypoxic conditions induced by cobalt chloride (CoCl_2_) treatment (100µM, Figure S2A) for 24 h, as confirmed by the protein expression of HIF-1α (Figure 5C). Nrf2-blockade was achieved by pre-treating differentiated cells with the Nrf2 suppressor brusatol (50nM, Figure S2B) for 3 h, and then the cells were subjected to CoCl_2_-induced hypoxia for 24 h (Figure 5D). Brusatol was in the medium until the experimental timeline ended. Protein levels associated with the Nrf2-ARE signalling pathway, iron metabolism, and ferritinophagy were assessed post-treatment. In agreement with *in vivo* ISO-induced MI, CoCl_2_ treatment increased the nuclear translocation of Nrf2, indicating enhanced Nrf2 transcriptional activity. This was further substantiated by the elevation in downstream targets, including GSR, NQO1, HMOX1, and GPX4 (P < 0.05). Conversely, Nrf2-blockade through brusatol resulted in decline in the levels of Nrf2 and subsequent downregulation of its downstream targets, suggesting the effective inhibition of Nrf2 signalling (Figure 5D, 5E). Our *in vitro* data conclusively demonstrated that Nrf2 suppression in cardiomyocytes reversed hypoxic stress, as corroborated by the HIF-1α levels (Figure 5D, 5E). Subsequently, we investigated changes in iron metabolism proteins following CoCl_2_ and brusatol administration. CoCl_2_ treatment elevated hepcidin levels, whereas brusatol effectively downregulated CoCl_2_-induced hepcidin levels (P < 0.05). Interestingly, consistent with the *in vivo* administration of ISO, no discernible changes were detected in FPN-1 levels following CoCl_2_ treatment. However, a significant augmentation (p < 0.05) in FPN-1 levels was observed upon brusatol treatment. Next, we performed a colorimetric analysis to determine iron levels using ferrozine assay (Figure 6A). In concordance with elevated Labile Iron Pools (LIPs) in ISO-induced MI, CoCl_2_-induced hypoxia led to increased labile iron levels, whereas brusatol administration significantly decreased labile iron levels (P < 0.05). Furthermore, we assessed whether suppression of Nrf2-transcriptional activity led to a decrease in ferritinophagic flux. Protein expression analysis of NCOA4, p62, Beclin-1, and LC3-I/LC3-II revealed that CoCl_2_-induced hypoxia increased ferritinophagy, whereas brusatol treatment reversed these effects (Figure 6B, 6C), potentially resulting in downregulation of iron sequestration in H9c2 cardiomyocytes.

**Figure 5:**
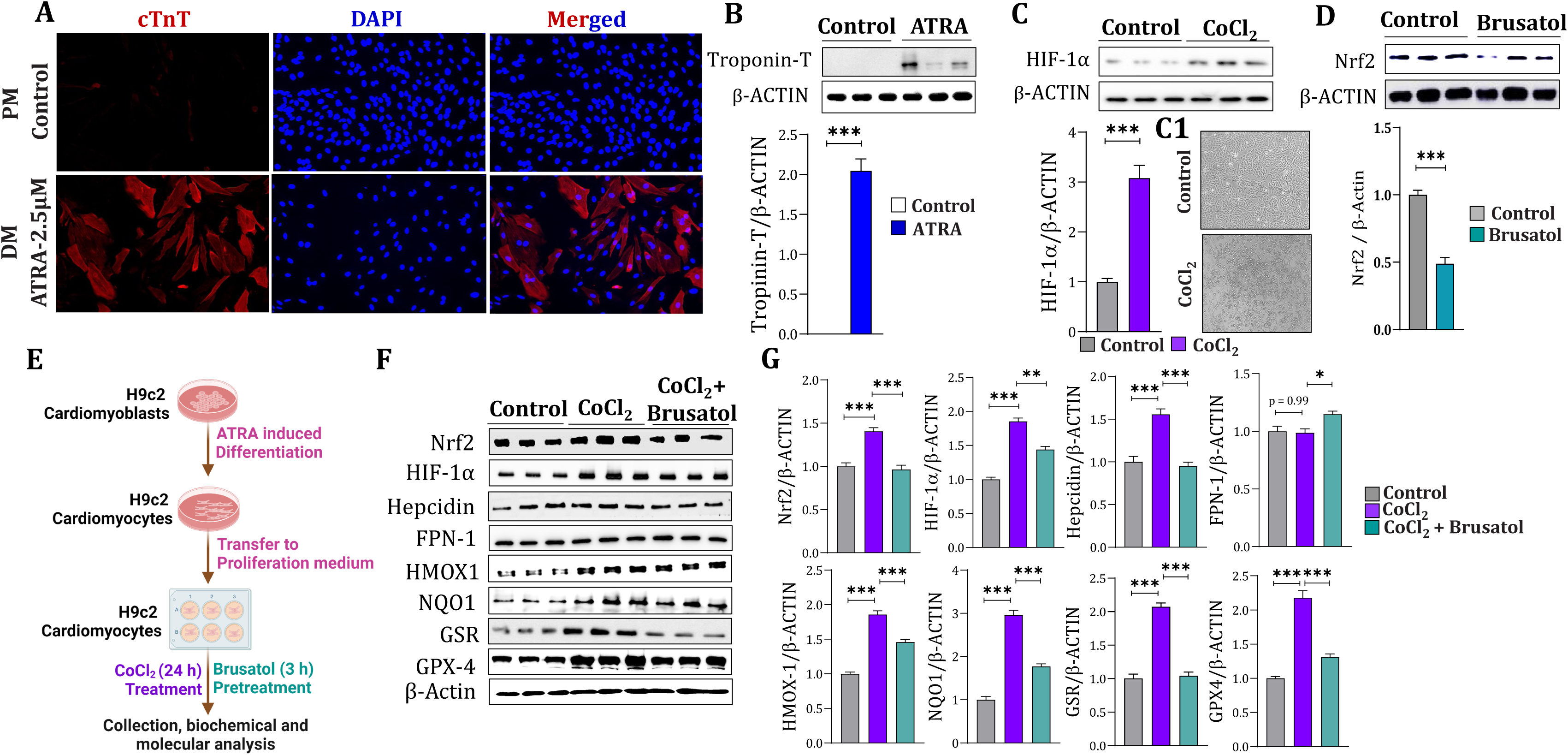
Pharmacological suppression of Nrf2 rescues hypoxic stress-induced changes in iron metabolism. H9c2 cardiomyoblasts were cultured in proliferation medium to 70% confluence and then subcultured for differentiation studies. Following 24 h of proliferation, the cells were grown in differentiation medium (DMEM comprising 0.5% FBS) containing all-trans-retinoic acid (ATRA, 2.5 μM) to induce differentiation up to 5 days. **(A)** Fluorescence images of cardiac troponin T staining in the control and ATRA-treated H9c2 cells. Following ATRA treatment, H9c2 cardiomyoblasts differentiated into cardiomyocytes, as indicated by the intense staining of cTnT. **(B)** Differentiation was further confirmed by immunoblotting of cTnT in control and ATRA-treated H9c2 cell lysates. **(C)** Immunoblot analysis of HIF-1α expression in H9c2 cells treated with 100 µM CoCl_2_ (n = 6). (C1) Bright-field images of H9c2 cardiomyocytes with or without CoCl_2_ treatment. **(D)** Immunoblot analysis of Nrf2 expression in H9c2 cells treated with 50 nM brusatol (n = 6). **(E)** Schematic depicting the experimental plan. Briefly, after ATRA-induced differentiation, H9c2 cardiomyocytes were transferred to a proliferation medium and pretreated with or without brusatol (50nM) for 3 h, followed by treatment with cobalt chloride (CoCl_2_, 100µM) for 24 h to induce hypoxic stress. **(F)** Immunoblot analysis of Nrf2, HIF-1α, hepcidin, FPN-1, HMOX-1, NQO1, GSR, and GPX4 expression in H9c2 cardiomyocytes treated with or without CoCl_2_ and brusatol. CoCl_2_ induced hypoxia leads to an increase in the protein levels of Nrf2, HIF-1α, hepcidin, HMOX1, NQO1, GSR, and GPX4, while there was no change in FPN-1. Interestingly, brusatol pretreatment reversed the effects of CoCl_2_, as evidenced by the suppression of Nrf2 levels and subsequent targets, including HIF-1α, hepcidin, HMOX1, NQO1, GSR, and GPX4. However, FPN-1 expression was significantly augmented by brusatol pretreatment. **(G)** Respective densitometries of Nrf2 (mean 1.00 vs 1.40 vs 0.96, p < 0.001), HIF-1α (mean 1.00 vs 1.85 vs 1.44, p < 0.001), Hepcidin (mean 1.00 vs 1.56 vs 0.95, p < 0.001), FPN-1 (mean 1.00 vs 0.99 vs 1.15, p = 0.99, p = 0.0198), HMOX1 (mean 1.00 vs 1.85 vs 1.46, p < 0.001), NQO1 (mean 1.00 vs 2.95 vs 1.77, p < 0.001), GSR (mean 1.00 vs 2.07 vs 1.04, p < 0.001) and GPX4 (mean 1.00 vs 2.18 vs 1.31, p < 0.001), n = 6 per group. All data are presented as the mean ± SEM. Statistical significance was calculated using one-way analysis of variance (ANOVA) (*p < 0.05, **p < 0.01, ***p < 0.001).

**Figure 6:**
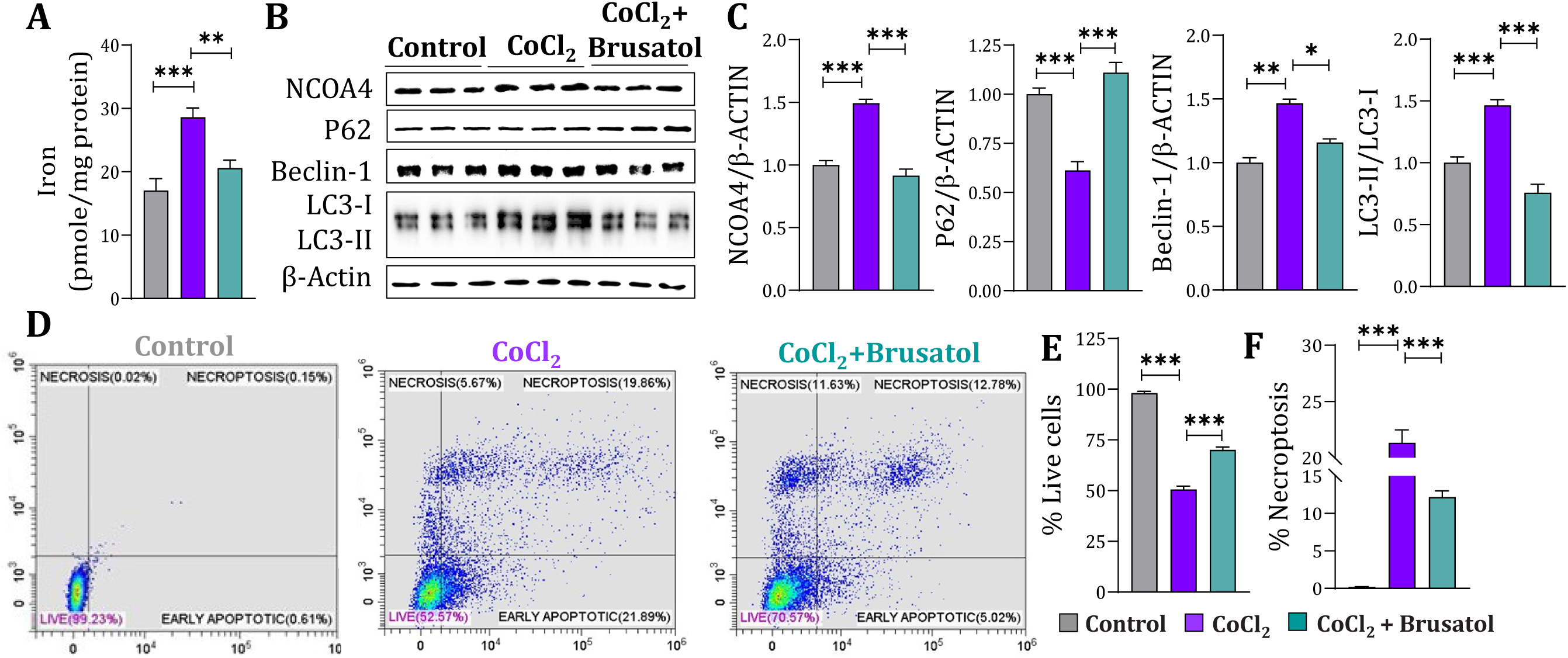
Pharmacological suppression of Nrf2 prevents ferritinophagy-induced cardiomyocyte death. **(A)** Bar graph depicting iron levels measured by colorimetric ferrozine assay in H9c2 cardiomyocytes treated with or without CoCl_2_ and brusatol. The results revealed a significant increase in the levels of iron upon CoCl_2_ induced hypoxia which was reversed by pretreatment with brusatol (mean 17.03 vs 28.61 vs 20.58, p = 0.0003, p = 0.0071), n = 6 per group. **(B)** Immunoblot analysis of NCOA4, P62, Beclin-1, LC3-I, and LC3-II levels in H9c2 cardiomyocytes treated with or without CoCl_2_ and brusatol. CoCl_2_ induced hypoxia leads to elevated levels of NCOA4, Beclin-1 and LC3-I/LC3-II, while P62 is downregulated. Pretreatment with brusatol abolished the changes induced by CoCl_2_**. (C)** Respective densitometries of NCOA4 (mean 1.00 vs 1.493 vs 0.92, p < 0.001), p62 (mean 1.00 vs 0.62 vs 1.11, p < 0.001), Beclin-1 (mean 1.00 vs 1.47 vs 1.16, p < 0.001), and LC3 (LC3-II/LC3-I ratio, mean 1.00 vs 1.47 vs 0.76 p < 0.001), n = 6 per group. **(D-F)** Flow cytometry analysis in H9c2 cardiomyocytes to determine the apoptosis after inducing hypoxia and brusatol administration. **(D)** Dot plot showing the extent of apoptotic cells in different treatment conditions, X-axis: Propidium iodide (PI); Y-axis: Annexin V FITC. **(E)** Bar graph showing the percentage of live cells (mean 98.10 vs 50.57 vs 70.07, p < 0.001) in different treatment conditions. **(F)** Bar graph showing the percentage of necroptosis (mean 0.19 vs 21.33 vs 12.17, p < 0.001) in different treatment conditions (n =5 per group). All data are presented as the mean ± SEM. Statistical significance was calculated using one-way analysis of variance (ANOVA) (*p < 0.05, **p < 0.01, ***p < 0.001).

**Figure 7:**
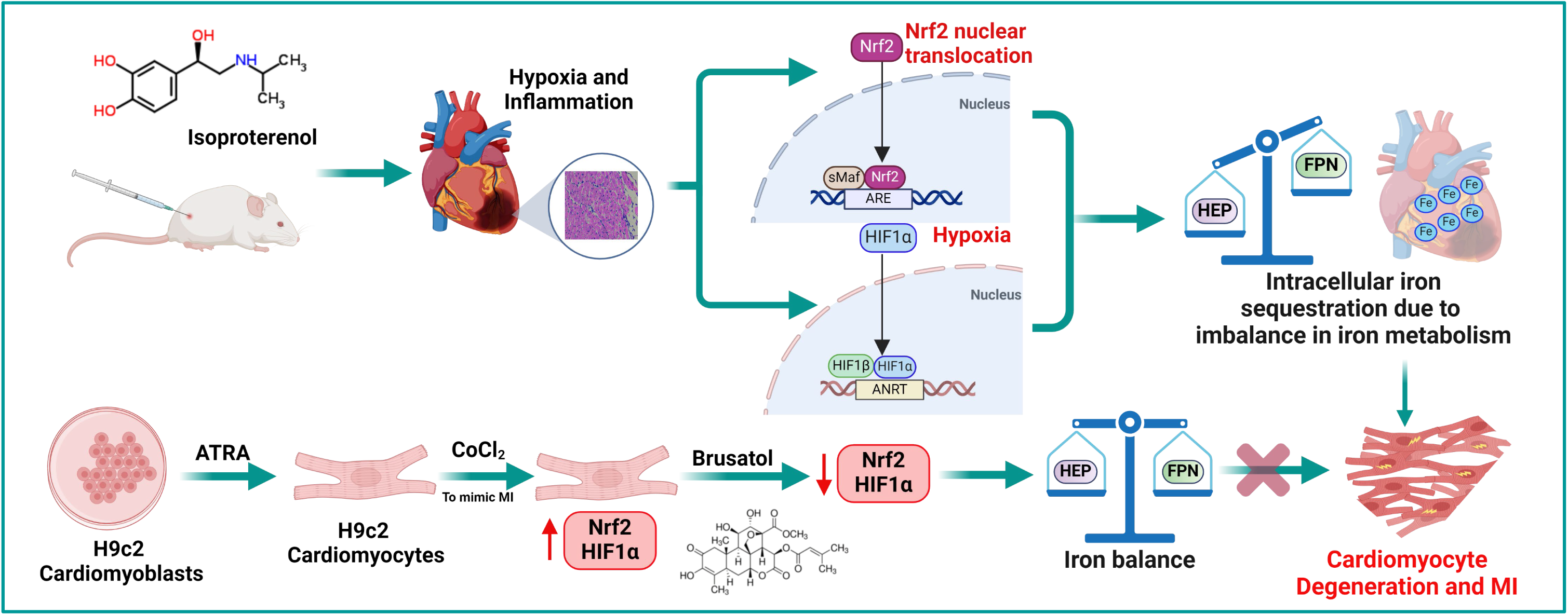
Schematic depicting the molecular mechanism underlying Nrf2 mediated MI pathogenesis through alterations in iron metabolism.

As an alternative measure of elevated ferritinophagy, we evaluated the apoptotic death rate after treatment with CoCl_2_ and brusatol. Annexin V staining followed by FACS demonstrated that CoCl_2_-induced hypoxia increased the number of apoptotic cells compared to the control, while brusatol treatment significantly (P < 0.001) mitigated apoptotic cell death (Figure 6D). In summary, our proof-of-concept experiments vividly demonstrate that MI/hypoxia induces alterations in iron metabolism-related proteins. Crucially, the hepcidin/ferroportin-1 ratio was elevated, leading to iron sequestration within cardiomyocytes. This, in turn, activates ferritinophagic flux, contributing to cardiomyocyte disintegration and death.

## 4. DISCUSSION

In redox biology, the maintenance of a stable intracellular redox network within the cellular environment has gained substantial attention over the past decade. The Nrf2/ARE transcription system has emerged as a focal point in maintaining redox balance and coordinating the anti-inflammatory response, suggested to protect against cardiovascular diseases, such as MI and atherosclerosis (Shen et al., 2019, Mimura & Itoh, 2015). Nonetheless, the exact impact of Nrf2 on the cardiovascular system remains uncertain due to conflicting findings in recent studies. As stated, Nrf2 has been reported to exhibit both beneficial and detrimental effects on myocardial infarction (MI) and other cardiovascular diseases animal models. (Ashrafian et al., 2012; Cao et al., 2015; Qin et al., 2016; Ma et al., 2019; da Costa et al., 2019; Zang et al., 2020; Li et al., 2022). Notably, our results highlighted, Nrf2 expression was augmented in the myocardial nucleus during ISO-induced MI. Nrf2 has a short half-life and is rapidly synthesized and degraded to maintain a stable equilibrium under homeostatic conditions. This resulted in minimal Nrf2 nuclear protein levels in control rats. However, under oxidative stress, Nrf2 degradation is impeded, leading to extended half-life, and increased nuclear levels. This phenomenon likely accounts for the significant increase in Nrf2 transcriptional activity observed in ISO-induced MI rats compared to that in control rats. Our results are in line with the findings of Li et al., 2022, which demonstrated Nrf2’s role in driving atherosclerosis and plaque complexity. Additionally, Mi et al., 2023 reported decreased Nrf2 protein levels in ISO-induced MI, conflicting with our experimental data. Given Nrf2’s pleiotropic nature, these discrepancies may stem from varied systemic and local effects and significant induction of the disease (Dodson et al., 2019b). In order to reconcile the contrasting outcomes and validate the heightened Nrf2 transcriptional activity, we assessed the levels of downstream targets, including NQO1, HMOX1, GSR, and GPX4. All of these indicators were found to be elevated in ISO-induced MI. However, enhanced lipid peroxidation was observed in ISO-induced MI, while there were no significant changes in protein carbonyls, adding complexity to the interpretation of the redox environment in ISO-induced MI. Upregulation of antioxidants appears to represent an adaptive cellular response aimed at maintaining redox homeostasis. While lipid peroxidation indicates oxidative stress, the activation of the Nrf2 system works to restore balance. Nevertheless, the Nrf2 response was not sufficient (at least in the tested acute phase, i.e. two days) to effectively counter the oxidative stress induced by ISO, resulting in the concurrent upregulation of both oxidants and antioxidants in the myocardium of ISO-administered animals.

Moreover, a growing body of evidence suggests that Nrf2 signalling contributes to initiating and sustaining the hypoxic response by influencing HIF-1α. Hence, we determined HIF-1α levels and found them to be augmented in ISO-induced myocardium compared to control rats. Previous studies have also reported elevated HIF-1α levels in ISO-induced MI (Jayachandran et al., 2010; Hosseinabadi et al., 2021; Hareeri et al., 2023). Multiple studies have shown that suppression of Nrf2 leads to a reduction in HIF-1α at the post-translational level (Toth & Warfel, 2017). Thus, augmented HIF-1α might be due to the hyperactive Nrf2 transcriptional activity observed in ISO-induced MI. HIF-1α mediates cardioprotection induced by ischemic preconditioning; however, prolonged hypoxic conditions can lead to aberrant cardiac remodeling and cardiac fibrosis. Fibrosis is characterized by excessive deposition of the extracellular matrix (ECM), including different types of collagens, hyaluronic acid, fibronectin, and proteoglycans in organs or tissues (Ho et al., 2014). In line with the elevated HIF-1α levels, we also observed enhanced collagen deposition through Masson’s trichrome staining upon ISO-induced MI. HIF-1α contributes to the upregulated gene expression of several ECM and non-ECM components, both *in vitro* and *in vivo* (Petrova et al., 2018, Dekker et al., 2022). At the same time, HIF-1α deficiency results in impaired collagen secretion in the presence of hypoxia (Hong et al., 2014; Xiong & Liu, 2017). Fibrosis is caused by inflammatory responses triggered by various stimuli like hypoxia and ischemic injury. During MI, the hypoxic milieu stimulates HIF-1α, which in turn activates neutrophils, enhancing their migration, phagocytic ability, and production of pro-inflammatory cytokines, resulting in inflammation and tissue damage. These findings indicate that ISO-induced MI leads to hypoxia, inflammation, and tissue damage with HIF-1α activation influenced directly by ISO and indirectly by increased Nrf2 transcriptional activity.

Pioneering studies have underscored the interplay between oxygen and iron regulation, demonstrating increased iron absorption in mice and rats when exposed to hypoxic conditions. (Renassia and Peyssonnaux 2019). These findings elucidate the intricate interconnections between oxygen homeostasis, erythropoiesis, and iron metabolism, tightly linking the physiology of the hypoxic response to the control of iron availability. Conversely, Nrf2 also reported to regulate genes associated with heme and nonheme-associated iron homeostasis (Campbell et al., 2013; Kerins & Ooi, 2018). Hence, we studied the intricate changes in iron metabolism during ISO-induced MI, on correlation with augmented Nrf2 and HIF-1α. Initially, we determined the labile iron pool in the myocardium and found that it was higher in ISO-induced rats than in the control rats. At gene level, both hepcidin and ferroportin-1 were observed to be elevated; however, only hepcidin followed the same trend at the protein level, whereas ferroportin-1 did not show any significant changes. This can be explained through two mechanisms: (1) Hepcidin regulates cellular iron efflux by binding to ferroportin and inducing its internalization. Post-translational regulation of ferroportin by hepcidin may thus complete a homeostatic loop that regulates the secretion of hepcidin, which in turn controls the level of ferroportin on the cell surface. Thus, sustained Nrf2 transcriptional activity during ISO-induced MI leads to steady-state levels of both hepcidin and ferroportin. However, post-translationally, as the iron content and hepcidin increase, ferroportin is rapidly degraded, balancing its level to no change upon ISO-induced MI. (2) Hypoxia significantly enhances hepcidin induction via STAT3 and Bmp6 signaling pathways (Silva et al., 2018; Jiang et al., 2023), a finding that aligns with our results showing elevated gene expression levels of Stat3, Bmp6, and Smad1 in ISO-induced myocardial infarction compared to the control group. Hypoxia-inducible factors, including HIF-1α and HIF-2α, have been reported to downregulate hepcidin (Liu et al., 2012); however, cardiac hepcidin levels rise in hypoxic conditions (Lakhal-Littleton et al., 2016). Furthermore, a recent clinical study identified that the increase in HIF-1α and HIF-2α does not seem to inhibit the increase in hepcidin levels in patients with chronic kidney disease (Hong et al., 2022). Similarly, in our study, it is possible that ISO-induced hypoxia led to hepcidin augmentation in an HIF-1α independent manner. Overall, these findings suggest that despite Nrf2’s role in inducing both hepcidin and ferroportin, hepcidin ultimately prevails as it is activated through multiple pathways and retains iron within myocardial cells during MI.

Numerous studies have demonstrated that ferritin (FTH and FTL) and ferroportin-1, crucial nonheme-associated iron regulatory proteins, are recognized transcriptional targets of Nrf2 (Wasserman and Fahl, 1997). At the gene level, our findings revealed an augmentation in both FTH and FTL, consistent with their status as transcription targets of Nrf2. However, at the protein level, FTL exhibited a similar trend, while FTH appeared to decrease twofold. One possible explanation for this would be; Ferritin L and H subunits are subject to distinct regulatory mechanisms both transcriptionally and post-transcriptionally. Moreover, FTH might function as a modulator of iron levels, responding to immediate demands of iron homeostasis rather than chronic overload or chelation (Cozzi et al., 2004). Interestingly, Sammarco et al. (2008) reported differential expression patterns of FTL and FTH under normoxic and hypoxic conditions following exposure to 50 ug/ml ferric ammonium citrate (FAC). Specifically, FTL-IRE levels demonstrated a twofold increase in HYP/FAC compared to NORM/FAC, whereas FTH IRE showed only a 25% increase in hypoxia. Consequently, FTL expression surpassed FTH during hypoxia due to differential regulation by IRP1 and IRP2, respectively. Similarly, in ISO-induced hypoxia, IRP1 and IRP2 might differentially regulate FTH and FTL post-transcriptionally. Furthermore, the concurrent increase in NCOA4 levels suggests rapid activation of ferritinophagy during ISO-induced MI. This in addition elucidates the protein-level downregulation of FTH, as NCOA4 facilitates FTH degradation and liberation of labile iron. Previous research has shown that isoproterenol increases the labile iron pool, induces ferritinophagy, and mediates cardiomyocyte death via an NCOA4-dependent pathway (Ito et al., 2021). Besides, ferritinophagy was confirmed by determining the downstream effectors of autophagy, including Beclin1, p62, and LC3-II/LC3-I. NCOA4-dependent ferritinophagy is regulated by intracellular iron (Dowdle et al., 2014; Mancias et al., 2014) and is associated with the Nrf2/HMOX-1 pathway in ferroptosis-induced cardiomyocyte death and heart failure (Dodson et al., 2019a; Li et al., 2020; Ito et al., 2021). Several studies have reported that Nrf2 prevents ferroptosis by increasing GPX4 and culminating in lipid peroxidation (Dodson et al., 2019a; Zhao et al., 2022). However, in our study, we observed elevated GPX4, NCOA4 and iron levels simultaneously. This maladaptive hyperactivation of ferritinophagy in MI hearts despite an abundant antioxidant response is a phenomenon that requires further investigation. However, NCOA4 expression is upregulated in an HIF-1α/HIF-2α-dependent manner under conditions that promote HIF stabilization (Li et al., 2020). Thus, upon ISO induction, HIF-1α stabilizes, leading to an increase in NCOA4, and consequently, FTH degradation. Additionally, sustained activation of Nrf2 leads to an enhanced antioxidant response, characterized by increased GPX4 and HMOX1 expression, and modifications in iron metabolism, as evidenced by a higher hepcidin/FPN-1 ratio. This leads to an increase in labile iron, which activates NCOA4-mediated ferritinophagy, regardless of Nrf2-mediated activation of GPX4. Recent findings by Federti et al. (2023) revealed a significant increase in non-transferrin bound iron and severe cardiac protein oxidation, both of which were linked to the activation of Nrf2 and the subsequent upregulation of related antioxidant and cytoprotective systems. Furthermore, the pressure overload-induced increase in GPX4 levels was compensatory rather than preventative to iron-dependent necrosis in cardiomyocytes (Ito et al., 2021). Thus, while GPX-4 activation is facilitated through Nrf2, NCOA4 activation occurs directly through HIF-1α and indirectly through Nrf2-mediated activation of hepcidin, resulting in dual nodes of ferritinophagy activation. Despite maintaining sustained Nrf2 transcriptional activity upon ISO treatment, NCOA4-mediated ferritinophagy could not be prevented.

To validate these findings *in vitro*, Nrf2 transcriptional activity was inhibited by using brusatol (Olayanju et al., 2015). Our findings demonstrate that administering brusatol to differentiated H9c2 cardiomyocytes under hypoxic conditions leads to a decrease in Nrf2 transcriptional activity. This decrease subsequently results in reduced expression of HIF-1α and hepcidin. Concurrently, there is an increase in the expression of ferroportin-1 (FPN-1), ultimately leading to a reduction in the labile iron pool. These findings contradict earlier research suggesting that Nrf2 plays a crucial role in regulating ferritinophagy/ferroptosis and in preventing cardiovascular diseases, including MI (Qin et al., 2021). However, it should be noted that clinical trials using antioxidants have failed, highlighting the need for a more precise understanding of the contribution of Nrf2-mediated antioxidant signalling in heart failure. Very recently, it has been demonstrated that there is a higher expression of Nrf2 in MI (Zuberi et al., 2024). Our study suggests that despite the active antioxidant system through Nrf2 during MI, the ferritinophagic flux is also activated and exacerbates the pathogenesis. Furthermore, our findings support the hypothesis that the inhibition of Nrf2/ARE signalling is preventative in ISO-induced MI or CoCl_2_-induced hypoxia.

## 5. CONCLUSION

In summary, the data presented herein elucidates pathological changes during myocardial infarction (MI) and the pivotal role of Nrf2/ARE signaling in MI pathology. Isoproterenol and/or CoCl_2_ induced significant alterations in redox signaling and hypoxia, leading to an increase in Nrf2/ARE signaling as a defensive response. While Nrf2 activation attempts to mitigate oxidative damage, it adversely affects iron metabolism. Both hepcidin and ferroportin are transcription targets of Nrf2 that play a key role in maintaining intracellular iron levels. Augmentation in Nrf2 leads to elevation in hepcidin; this hepcidin-ferroportin imbalance triggers ferritinophagy and subsequent increase of labile iron in cardiomyocytes under isoproterenol or CoCl2 stress. Notably, brusatol pretreatment mitigates this imbalance and prevents ferritinophagy, thereby preventing cardiomyocyte cell death.

## 6. LIMITATIONS AND FUTURE PERSPECTIVES

This study has certain limitations. Despite our efforts to explore the influence of the Nrf2/ARE signaling pathway on myocardial infarction (MI), we recognize that utilizing transgenic models with either Nrf2 overexpression or knockout would offer clearer insights into this crucial signaling cascade. Furthermore, employing whole-genome or single-cell sequencing specific to cardiomyocytes would aid in understanding Nrf2/ARE signaling and its role in fibrosis, inflammation, iron metabolism, and ferritinophagy under both normal and disease conditions, such as MI. Hence, our future endeavors include investigating the transcriptomic alterations during MI in models where Nrf2 is manipulated. Additionally, the observed disparities in ferritin subunits require validation through elucidating post-transcriptional mechanisms in both naïve and MI conditions. Intriguingly, there is limited understanding of the serum hepcidin/ferritin ratio in the South Indian population. Therefore, a clinical investigation is warranted to discern alterations in hepcidin, ferritin, and other iron-related parameters across various cardiovascular diseases in South Indian population. Ultimately, the present study and our future prospects aim to provide insights into the intricate interactions governing the redox environment, oxygen demand, and iron metabolism in both normal and diseased conditions.

## Supporting information

Key resources table

## AUTHOR CONTRIBUTIONS

***Deepthy Jayakumar***: Conceptualization (First); data curation (First); formal analysis (First); investigation (First); methodology (First); visualization (First); writing − original draft (First); writing − review and editing (First). ***Kishore Kumar S. Narasimhan***: Conceptualization (supporting); formal analysis (supporting); visualization (supporting); writing − original draft (supporting); writing − review and editing (supporting). ***Abinayaa Rajkumar***: Investigation and visualization (supporting). ***Gokulprasanth Panchalingam***: Investigation and visualization (supporting). ***Navvi Chandrasekar***: Investigation and visualization (supporting). ***Varsha C. Ravikumar***: Investigation and visualization (supporting). ***Kalaiselvi Periandavan***: Conceptualization (lead); funding acquisition (lead); project administration (lead); resources (lead); supervision (lead); validation (lead); visualization (lead); writing − review and editing (lead). All authors have read and approved the final version of the manuscript.

## ACKNOWLEDGEMENTS

Financial assistance to Deepthy Jayakumar, Department of Medical Biochemistry, from the Council of Scientific and Industrial Research (CSIR), in the form of the CSIR Junior Research Fellowship (JRF) and Senior Research Fellowship (SRF), New Delhi, Government of India, is gratefully acknowledged. The authors extend their gratitude to Dr. Priya Ramanathan, Associate Professor, and Ms. Anusha Jayachander from the Department of Molecular Oncology at the Cancer Institute Adyar (WIA) for their invaluable technical assistance in performing and analyzing Flow Cytometry. The authors express their gratitude to the Department of Health Research - Multidisciplinary Research Unit (DHR-MRU) at the Dr. ALM Post Graduate Institute for Basic Medical Sciences, University of Madras, for providing access to its state-of-the-art facilities.

## DATA AVAILABILITY STATEMENT

The data supporting the findings of this study are available from the corresponding author upon reasonable request.

## DECLARATION OF TRANSPARENCY AND SCIENTIFIC RIGOUR

This Declaration acknowledges that this paper adheres to the principles for transparent reporting and scientific rigor of preclinical research recommended by funding agencies, publishers, and other organizations engaged in supporting research.

## Supplemental Figure Legends

**Figure S1.**
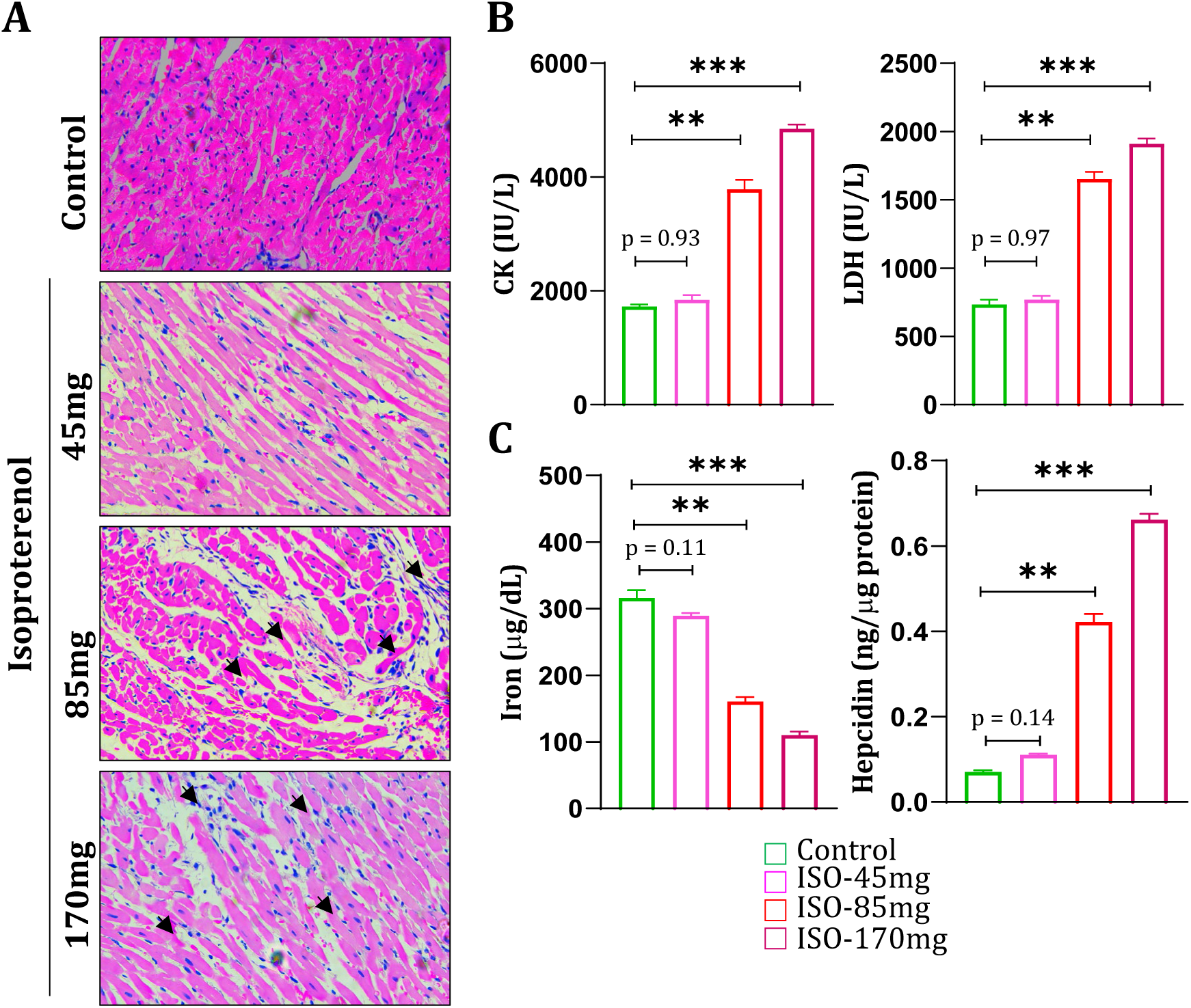
Dose-dependent study to determine the optimal ISO dose to induce MI. Briefly, 45, 85, and 170 mg/kg ISO were administered subcutaneously to the rats for two days. **(A)** Paraffin sections show inflammatory infiltration (black arrows) in ISO 85 and 170 myocardium compared to control by H&E staining. Furthermore, more necrotic impressions were observed in the animals treated with 170 mg, while 45 mg did not show any significant pathological hallmarks of MI. **(B)** Bar graphs depicting the levels of creatine kinase and lactate dehydrogenase in the serum of control ISO (45 mg, 85 mg, and 170 mg) animals. **(C)** Bar graphs depicting the levels of iron and hepcidin in the serum of control ISO (45 mg, 85 mg, and 170 mg) animals. All data are presented as the mean ± SEM. Statistical significance was calculated using one-way analysis of variance (ANOVA) (*p < 0.05, **p < 0.01, ***p < 0.001).

**Figure S2:**
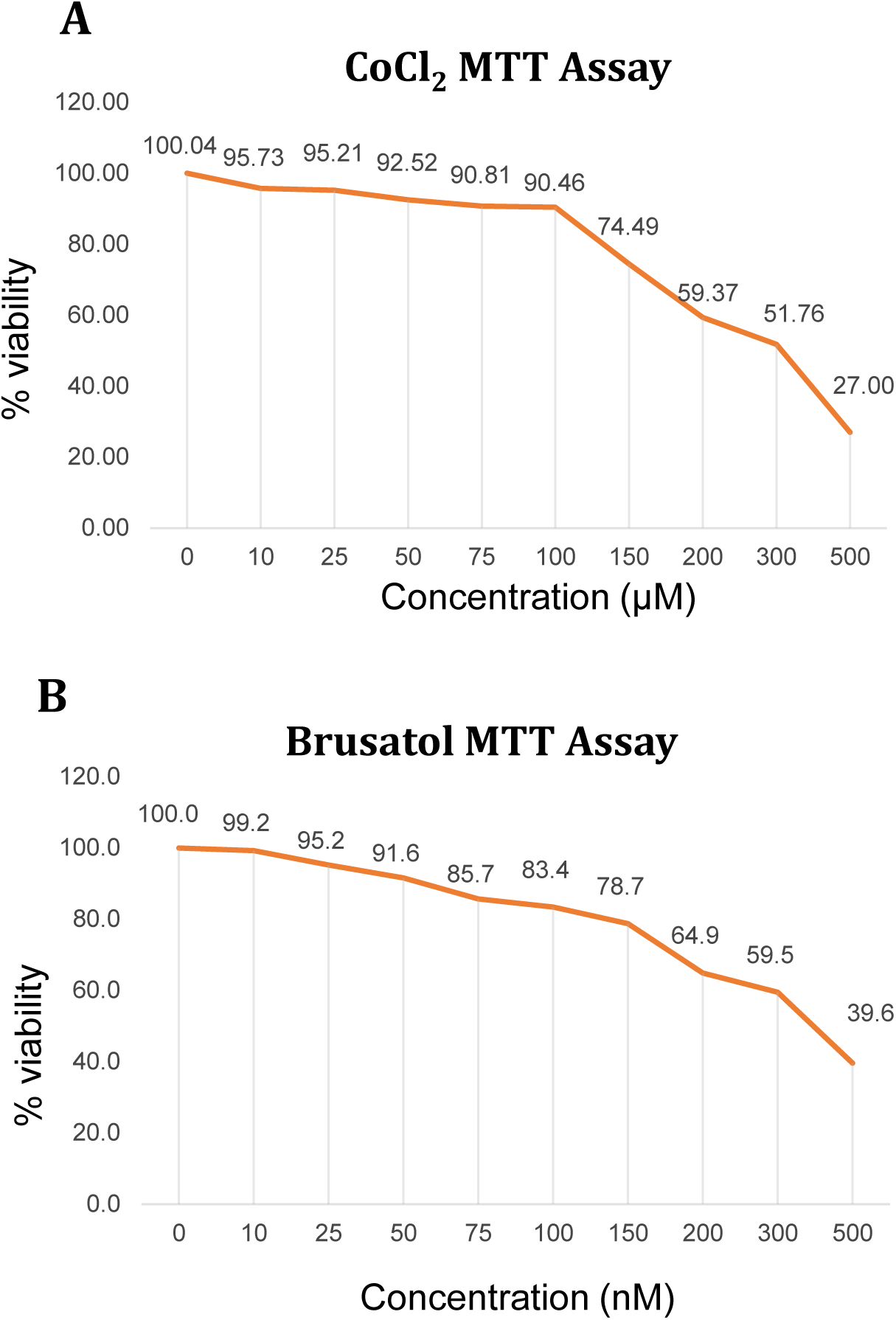
Determination of the optimal doses of CoCl_2_ and brusatol. **(A)** Graphical representation of MTT readouts of H9c2 cells treated with different concentrations of CoCl_2_, ranging from 0 to 500 µM. When treated with CoCl_2_ at 100 µM, 90.81% of the cells were viable; this dose was chosen as the optimal concentration. **(B)** Graphical representation of the MTT readouts of H9c2 cells treated with different concentrations of brusatol (0–500 nM). When treated with 50 nM brusatol, 91.6% of the cells were viable, and this dose was chosen as the optimal concentration.

